# Multi-omics reveals global signaling rewiring and identifies Activin A-induced dysregulation of FOS/Activator Protein 1 as a novel target in Fibrodysplasia ossificans progressiva

**DOI:** 10.1101/2025.01.21.634061

**Authors:** Marius Wits, Nerea Gómez-Suárez, Alexandre Deber López, Nicole Farfán, Fjodor Bekedam, Bharath Sampadi, Sarah Rotman, Daniel Rubiera López, Judit Bestilleiro Márquez, Christiaan Arendzen, Christian Freund, Peter van Veelen, Aida Llucià Valldeperas, Frances de Man, Francesc Ventura, Marie-José Goumans, Gonzalo Sanchez-Duffhues

**Affiliations:** Department of Cell and Chemical Biology, Leiden University Medical Center, Einthovenweg 20, 2333 ZC, Leiden, the Netherlands; Nanomaterials and Nanotechnology Research Center (CINN), Spanish National Research Council (CSIC), Health Research Institute of Asturias (ISPA), 33011 Oviedo, Asturias, Spain; Departament de Ciències Fisiològiques, Universitat de Barcelona, IDIBELL, C/ Feixa Llarga s/n 08907 Hospitalet de Llobregat, Spain; PHEniX laboratory, Department of Pulmonary Medicine, Amsterdam UMC location Vrije Universiteit, Amsterdam, The Netherlands, Amsterdam Cardiovascular Sciences, Pulmonary Hypertension and Thrombosis, Amsterdam, The Netherlands; Center for Proteomics and Metabolomics (CPM), Leiden University Medical Center, Einthovenweg 10, 2333 ZC, Leiden, The Netherlands; Leiden hiPSC Center, Leiden University Medical Center, Silviusweg 62, 2333 BE, Leiden, The Netherlands

**Author notes:** Address for correspondence: Gonzalo Sanchez-Duffhues. These authors contributed equally (Marie-José Goumans, Gonzalo Sanchez-Duffhues).

**Keywords:** Phosphoproteomics, transcriptomics, Activin A, ACVR1, heterotopic ossification, mechanotransduction, BMP, FOS, JUN

## Abstract

**Background:** Fibrodysplasia ossificans progressiva (FOP) is caused by an activating mutation (p.R206H) in the type I BMP receptor ALK2, leading to heterotopic ossification (HO) in soft connective tissues. While aberrant Activin A-induced SMAD signaling is central in FOP pathogenesis, global signaling alterations remain poorly understood.

**Methods:** We performed phosphoproteomics, transcriptomics and biochemical analyses in mesenchymal cells (MSCs) overexpressing wild-type ALK2^WT^ or mutant ALK2^R206H^ receptors and in induced-MSCs derived from FOP patient iPSCs. Findings were validated in vivo using FOP-like mouse models and in vitro via pharmacological interventions.

**Results:** Multi-omics analyses revealed previously unrecognized signaling networks in ALK2^R206H^ cells, including enhanced MAPK, mTOR, RUNX2 and RHO-mediated mechanotransduction pathways. Notably, we identified dysregulated Activator Protein-1 (AP-1) expression and function as a novel contributor to FOP. AP-1 factors were highly enriched in HO lesions in FOP-like animals. Pharmacological inhibition of AP-1 significantly reduced osteochondrogenic differentiation in vitro.

**Conclusion:** This study highlights global signaling dysregulation in FOP and identifies AP-1 as a critical driver and potential therapeutic target for FOP.

## 1. Introduction

Fibrodysplasia ossificans progressiva (FOP) is an ultra-rare genetic disorder with an estimated prevalence of one in a million, characterized by heterotopic ossification (HO) in soft connective tissues including skeletal muscle.^1^ HO is triggered by spontaneous or injury-induced episodic inflammations, resulting in painful soft tissue swellings, referred to as flare-ups. Starting at pediatric age, cumulative formation of extra-skeletal bone leads to severe immobility and significantly impacts quality of life. The median age of mortality is 56 years, with cardiorespiratory failure caused by thoracic insufficiency syndrome being the most common cause of death.^2^

Conventional therapies for FOP are palliative, focusing on managing inflammation during the episodic flare-ups with treatments such as high-dose corticosteroids. However, no definitive cure exists today. Recently, the FDA has approved Sohonos (i.e. Palovarotene), a retinoic acid receptor-gamma agonist, for FOP patients from 8 (female) or 10 (male) years and older in the USA and Canada.^3^ Palovarotene reduces the amount of new HO formation, but is associated with substantial side-effects.^3^ While this approval represents a promising step forward, more efficient and safe therapies are urgently needed to treat or cure FOP.

Nearly all FOP patients harbor the gain-of-function mutation *ACVR1* c.617G>A, which encodes the mutant bone morphogenetic protein (BMP) type I receptor ALK2 (p.R206H).^4,5^ BMP ligands signal by binding to their dedicated serine/threonine receptors on the plasma membrane.^6^ BMPs are part of the structurally similar transforming growth factor-β (TGF-β) signaling family,^7^ whose classical downstream effectors are the small mothers against decapentaplegic (SMAD) proteins. The TGF-β signaling family exert pleiotropic effects and maintaining balanced signaling is crucial for fundamental cellular processes and tissue homeostasis.^8^

High affinity ligands for ALK2 are BMP6, BMP7 and Activin A. While BMP6 and BMP7 activate ALK2, Activin A acts as an ALK2 antagonist. However, in FOP, the mutant ALK2^R206H^ receptor becomes responsive to Activin A.^9^ Multiple studies have shown that the mutation renders the receptor unable to inactivate entirely, resulting in so-called leaky signaling activation.^10^ Clustering of ALK2^R206H^ receptors, whether induced by ligands such as Activin A, optogenetic activation^10^ or exogenous monoclonal antibodies targeting ALK2, is sufficient to initiate downstream signal transduction and HO development.^11^ Activin A is a pro-inflammatory factor produced by mesenchymal, chondrogenic cells (including fibroadipose progenitors) and immune cells including macrophages.^12,13^ Given the robust and potent inhibitory effect of an anti-Activin antibody in human clinical trials, it seems that this disturbed Activin A-induced ALK2^R206H^ signaling is a primary driver of HO in FOP.^14,15^ In addition to targeting canonical ALK2 and Activin A signaling, therapeutic approaches aiming at normalizing non-SMAD signaling have shown promising preclinical results.^16–18^ BMP receptors can act as serine/threonine and tyrosine kinases, mediating non-canonical pathways such as extracellular signal-regulated kinases (ERK), phosphoinositol-3 kinase (PI3K) or the p38 MAPK pathway.^19^ While canonical SMAD-mediated signaling downstream of Activin A – ALK2^R206H^ stimulation has been extensively studied, non-Smad signaling pathways remain poorly understood. We hypothesized that the p.R206H mutation in ALK2 might alter non-SMAD signaling networks, unveiling novel therapeutic targets. To test this hypothesis, we performed a multi-omics approach, including phosphoproteomics, transcriptomics and biochemical assays, on ALK2^R206H^-expressing hMSCs and FOP-patient derived iPSC induced-MSCs. These analyses revealed previously unrecognized signaling pathways with significant therapeutic potential for FOP.

## 2. Materials and Methods

### Generation of overexpression and isogenic cell lines

Human bone marrow-derived Mesenchymal Stem Cells (hMSCs) were transduced by lentiviral delivery encoding the ALK2/ACVR1^WT^ or ALK2/ACVR1^R206H^ receptors as described before.^16^ In short, the receptor constructs were subcloned into lentiviral plasmids whereafter 3^rd^ generation lentiviral particles were produced in HEK293t cells and the hMSCs were transduced.

The isogenic iPSCs (iso01LUMC0084iFOP02) were generated by the LUMC hiPSC Centre using Cas9- mediated gene editing of the ACVR1 c.617G>A mutation in the FOP patient-derived iPSCs line (LUMC0084iFOP02)^20^ directed by the crRNA sequence 5’-GCTCACCAGATTACACTGT-3’. Flow cytometry confirmed pluripotency following NANOG, SSEA4 and OCT3/4 staining after gene editing. In addition, in vitro trilineage differentiation with subsequent staining of the ectoderm markers Nestin, PAX6, FABP7, the endoderm markers Eomes, FOXA2, GATA4, and the mesoderm markers Vimentin, CDX2 and Brachyury further confirmed pluripotency of the isogenic FOP lineage.

### Cell culture and iMSC differentiation

HEK293t and COS-1 cells were cultured in DMEM (Gibco, cat. 11965092, Waltham, Massachusetts, USA) supplemented with 10% Fetal Bovine Serum (FBS) (Biowest, cat. S1810-500, Bradenton, Florida, USA) and 100 U/ml P/S. iPSCs were cultured on vitronectin-XF (STEMCELL Technologies, cat. 07180, Vancouver, Canada) coated plates with mTeSR-Plus medium (STEMCELL Technologies, cat. 100-0276, Vancouver, Canada). Differentiation of iPSCs towards an induced MSC (iMSCs) lineage was done using the STEMdiff^TM^ Mesenchymal Progenitor Kit (STEMCELL Technologies, cat. 05240) following manufacturer’s protocol. During iMSC differentiation the cells were dissociated with TrypLE Select 10x. hMSCs and differentiated iMSCs were cultured in Alpha-MEM (Gibco, cat. 32561094) supplemented with 10% FBS, 1 ng/ml bFGF (Sigma-Aldrich, cat. F0291, St. Louis, Missouri, USA) and 100 U/ml P/S (growth medium; GM). All lines were cultured in a CO_2_ controlled 37°C incubator. Standard passaging was performed using Trypsin/EDTA incubation.

In the case of ligand stimulations, cells were serum deprived for 16 hours overnight using DMEM without supplements. Stimulation was performed using medium refreshment with 50 ng/ml Activin A (R&D, cat. 338-AC-010/CF, Minneapolis, Minnesota, USA) or BMP6 (R&D, cat. 507-BP-020/CF) with ligand buffer vehicle (4 mM HCl in 0.1% BSA) as control.

### Phosphoproteomics

#### Sample pretreatment

Cell lysis and digestion were performed as previously described.^21^ Cells were lysed using 5% sodium dodecyl sulfate (SDS) lysis buffer (100 mM TRIS-HCl pH 7.6) and incubated at 95 °C for 4 minutes. A Pierce BCA Gold protein assay (Thermo Fisher Scientific) was performed in duplicate to quantify the protein amount. 300 µg Protein of each sample was then reduced with 5 mM tris(2- carboxyethyl)phosphinehydrochloride (TCEP), followed by alkylation using 15 mM iodoacetamide (IAA). Excess IAA was quenched using 10 mM dithiothreitol (DTT). Protein lysates were precipitated using chloroform/methanol/water (4:1:3 v/v/v), then washed with methanol and air dried. The resulting protein pellets were re-solubilized in 40 mM ammonium bicarbonate pH 8.4 (ABC) and digested overnight at 37 °C with a trypsin/lysyl endopeptidase (LysC) mix (mass spec grade Trypsin/LysC mix, Promega) with an enzyme/substrate ratio of 1:60. Peptide quantification was performed in duplicate using the Pierce BCA Gold protein assay. 230 µg Peptide of each sample was then taken and desalted using 30 mg 1cc C18 SPE cartridges (Oasis HLB, Waters). The samples were washed with 0.1% trifluoroacetic acid (TFA), eluted with 80% acetonitrile (ACN)/0.1% TFA and lyophilized to subject the samples either to phosphopeptide enrichment or to store the samples at -20 °C until further use.

Automated phosphopeptide enrichment was performed through the AssayMap Bravo Platform (Agilent Technologies) following the Phosphopeptide Enrichment v2.1 protocol. Phosphorylated peptides were enriched using Fe(III)-NTA 5 µL cartridges (Agilent Technologies). The cartridges were primed with 250 µL of 99.9% ACN/0.1% TFA (v/v) at a flow rate of 300 µL/min and equilibrated with 250 µL loading buffer of 80% ACN/0.1% TFA at 5 µL/min. Samples were dissolved in 203 µL loading buffer, while 200 µL was loaded onto the cartridges at 10 µL/min. The columns were then washed with 250 µL loading buffer at 10 µL/min, and the phosphorylated peptides were eluted with 125 µL of 1% ammonia at 5 µL/min. Samples were lyophilized and stored at -20 °C until subjected to LC-MS/MS.

#### Liquid chromatography-mass spectrometry (LC-MS)

Peptides were dissolved in water/formic acid (100/0.1 v/v) and analyzed by on-line C18 nanoHPLC MS/MS with a system consisting of an Ultimate3000nano gradient HPLC system (Thermo, Bremen, Germany), and an Exploris480 mass spectrometer (Thermo). Samples were injected onto a cartridge precolumn (300 μm × 5 mm, C18 PepMap, 5 um, 100 A, and eluted via a homemade analytical nano-HPLC column (50 cm × 75 μm; Reprosil-Pur C18-AQ 1.9 um, 120 A (Dr. Maisch, Ammerbuch, Germany). The gradient was run from 2% to 40% solvent B (20/80/0.1 water/acetonitrile/formic acid (FA) v/v) in 120 min at 250 nl/min. The nano-HPLC column was drawn to a tip of ∼10 μm and acted as the electrospray needle of the MS source. The mass spectrometer was operated in data-dependent MS/MS mode, with a HCD collision energy at 30% and recording of the MS2 spectrum in the orbitrap, with a quadrupole isolation width of 1.2 Da. In the master scan (MS1) the resolution was 120,000, the scan range 400-1500, at standard AGC target and a maximum fill time of 50 ms. A lock mass correction on the background ion m/z=445.12003 was used. Precursors were dynamically excluded after n=1 with an exclusion duration of 45 s, and with a precursor range of 20 ppm. Included charge states were 2-5. For MS2 the first mass was set to 120 Da, and the MS2 scan resolution was 30,000 at an AGC target of 75% at a maximum fill time of 60 ms. In a post-analysis process, raw data were first converted to peak lists using Proteome Discoverer version 2.5 (Thermo Scientific), and then submitted to the minimal human Uniprot database (20596 entries), using Mascot v. 2.2.07 (www.matrixscience.com) for protein identification. Mascot searches were done with 10 ppm and 0.02 Da deviation for precursor and fragment mass, respectively, and trypsin was specified as the enzyme. Methionine oxidation and the acetylation (on the protein N-terminus) were set as variable modifications. Carbamidomethyl was set as a fixed modification on cysteines. The false discovery rate was set < 1%. Alternatively, Maxquant version 2.5.1.0 was used with default settings.

#### Data analysis

The raw data was processed using MaxQuant (v2.5.1) and mapped to the *Homo sapiens* UniProt proteome database (2023). Phospho(STY) was added to variable modifications. Digestion mode was set to specific with enzymes Trypsin/P and LysC/P. Maximum missed cleavages allowed was set to 4. Phosphosite intensity values were loaded in Perseus (v1.6.14.0), and log2 transformed. Further cleanup was performed to remove potential contaminants and reverse hits. Phosphosite table was further expanded using an inbuilt function and further filtered to retain quantifications in at least three replicates in any condition and with a localization probability of >0.75. A downshifted normal distribution was used to sample for missing value imputation with the following settings: N(width:0.3x σ, downshift:1.8x σ) where σ is the standard deviation of the measured values.

Significantly responsive phosphosites were identified by performing two-sample student’s t-test with a p-value < 0.05 and sufficient magnitude (1.5 fold-change) cut-off. For pathway enrichment analyses the differential phosphorylated protein was used irrespective of its phosphorylation site. Furthermore, the STRING environment (v12.0; string-db.org)^22^ was used for protein-protein interaction analyses, co-expression analyses and enrichment analyses including Reactome pathway analysis, all performed in October 2024. Separately, gene ontology enrichment analysis (GO Ontology database DOI: 10.5281/zenodo.12173881, released 2024-06-17; geneontology.org)^23,24^ was performed for additional tests. Upon enrichment analyses, the data was extracted accordingly. The generation of descriptive data and pathway analysis figures was performed using R (version 4.4.0) within the R Studio environment (version 2024.04.0), using the ’tidyverse’ (v1.3.0) and ’ggplot2’ (v3.5.1) packages.

### Bulk RNA sequencing

Human MSCs overexpressing ALK2^WT^ or ALK2^R206H^ were serum deprived for 16 hours following 1 hour stimulation of Activin A (50 ng/ml) or BMP6 (50 ng/ml). RNA samples were obtained using the ReliaPrep RNA Miniprep Systems (Promega, cat. Z6012, Madison, Wisconsin, USA) kit and subsequently sequenced using the Illumina NovaSeq 6000 (Illumina, San Diego, California, USA) platform through Novogene’s pipeline. Read mapping was performed using the Hisat2 (v2.0.5) package and the total/normalized counts were measured using featureCounts (v1.5.0-p3). Differential expression analysis (including principal component analyses) was performed using the R package DESeq2 (v1.20.0) following the Benjamini and Hochberg adjusted approach. Gene hits with an adjusted p-value < 0.05 were considered differentially expressed. Heatmaps and volcano plots were generated using the R packages ‘ComplexHeatmap’ (v2.20.0) and ‘tidyverse’ (v2.0.0) including ggplot2 (v3.5.1), respectively. RStudio was used to generate and document the RNA sequencing analyses. The whole dataset is publicly available (see GSE237512).

### RT-qPCR

RNA was isolated using the Reliaprep RNA Miniprep Systems (Promega, cat. Z6012) kit and cDNA was synthesized using the RevertAid First Strand cDNA Synthesis Kit (Promega, cat. K1621) following manufacturer’s protocol scaled to 0.5 µg RNA input. RT-qPCR was conducted in a 384-wells format (Bio-Rad, cat. HSP3801, Hercules, California, USA) of 8 µL total PCR reaction consisting of 2 µL 10x diluted cDNA, 2 µL 10 µM forward and reverse primer mix (custom from IDT, Leuven, Belgium), and 4 µL 2x GoTaq qPCR Master Mix (Promega, cat. A600X). The Bio-Rad CFX384 (Bio-Rad) was used to run the qPCR plates. The data was analyzed following an in-house analysis pipeline utilizing the standard double delta (Δ) Ct method with extra internal controls, subsequently, the fold change was shown by 2^-ΔΔCt and statistics was performed on the ΔΔCt. All used primer sets are depicted in the following table.

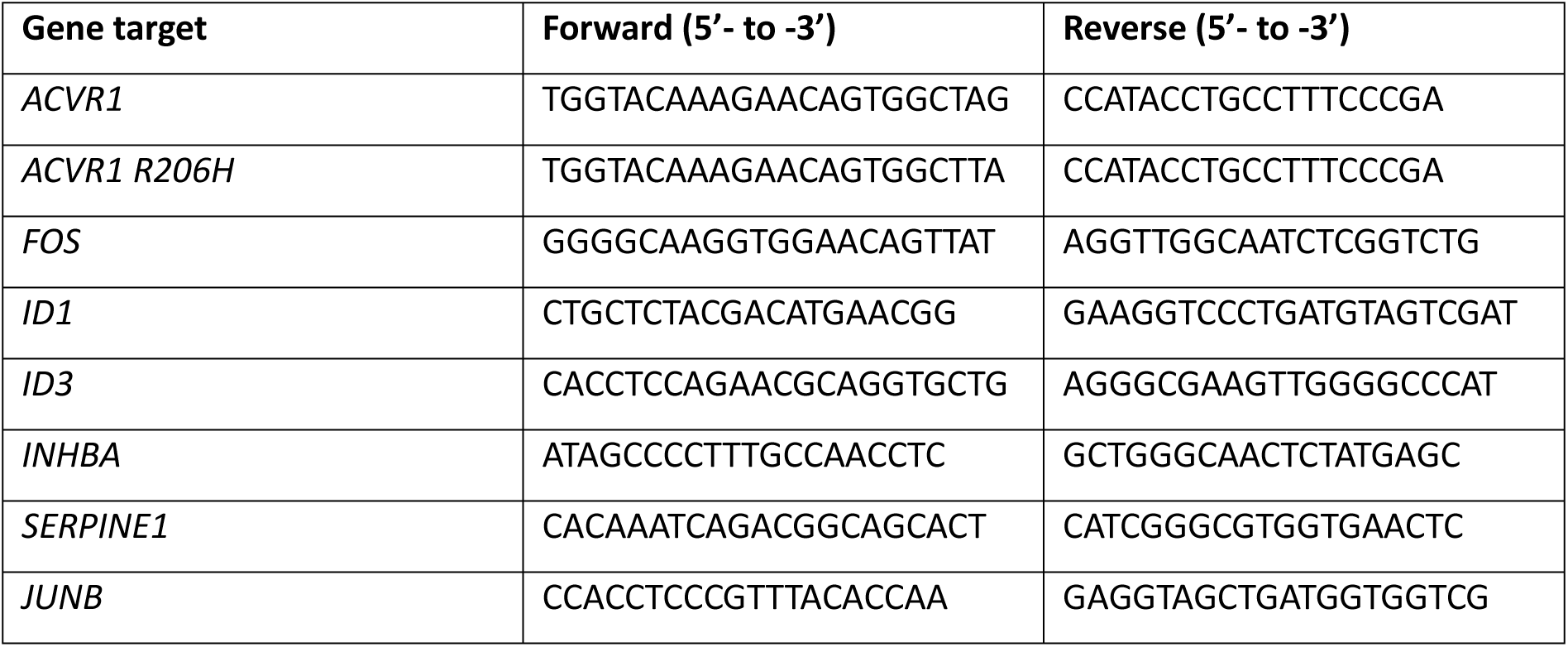

### SDS-PAGE and Western blotting

Cells were lysed using RIPA buffer and quantified using the Pierce BCA Protein assay (Thermo Scientific, cat. 23225) following standard protocols. The proteins were denatured using βME containing Laemmli buffer and equally loaded on a 10% polyacrylamide gel prior to SDS-PAGE. The separated proteins were blotted on PVDF membranes, blocked using 10% milk (Campina, Amersfoort, The Netherlands) in TBST and incubated with the primary antibody (1:1000 dilution) in 5% BSA (Sigma, cat. 05479) in TBST overnight at 4°C. The primary antibodies used were pSMAD1/5 (Cell Signaling, cat. 9515), SMAD1 (Cell Signaling, cat. 6944), pSMAD2 (home-made)^25^, SMAD2 (BD Biosciences, cat. 610842), or Vinculin (Sigma, cat. V9131). After thorough washing with TBST the membranes were incubated with the secondary antibody (1:10000, Anti Mouse-HRP, Promega, cat. W4021 or Anti Rabbit-HRP, Invitrogen, cat. 31458) in 10% milk in TBST for 2 hours at 4°C. The membranes were washed, ECL activated using WesternBright (Isogen, cat. K12042D20, Utrecht, The Netherlands), and imaged using the ChemiDoc (Bio-Rad).

### Luciferase reporter assays

Confluent 24-wells plates with COS-1 cells were transfected with ALK2^WT^ or ALK2^R206H^ plasmids (200 ng/well) with the MMP1- or mutated TRE MMP1-Luciferase reporter construct^26^ (100 ng/well), using a 1:2 DNA:PEI ratio. Similarly, confluent 24-wells plates of hMSC stably expressing ALK2^WT^ or ALK2^R206H^ were transfected with MMP1- or mutated TRE MMP1-Luciferase reporter constructs^27^ (100 ng/well) with a LACZ expression plasmid (100 ng/well) using Dharmafect DUO (Horizon Discovery, cat. T-2010- 02, Cambridge, UK). After overnight transfection, the medium was refreshed to the indicated conditions: vehicle, Activin A (100 ng/ml), or phorbol 12-myristate 13-acetate (PMA) (50 ng/ml) for 2 days. Subsequently, cells were washed with cold PBS and lysed with luciferase lysis buffer for 30 minutes on ice. Next, luciferase was measured using the Luciferase Assay System (Promega, cat. E1501) at 560 nm emission following manufacturer’s protocol. Β-galactosidase (βGal) activity was measured to control for transfection efficiency.

### Immunofluorescence and Masson’s trichrome staining

Using 8-well chamber slides (Thermo Fisher, cat. 154534), confluent hMSC-ALK2^WT^ or -ALK2^R206H^ cells were serum deprived for 16 hours and stimulated for 45 minutes with Activin A (100 ng/ml) or PMA (50 ng/ml). After washing with PBS the cells were fixated using 4% PFA in PBS at room temperature (RT) for 30 minutes and the aldehyde groups were quenched with 2 mg/ml glycine in PBS for 5 minutes. The samples were permeabilized using 0.2% Triton X-100 in PBS for 10 minutes at RT. After washing with PBS the samples were blocked for 1 hour using 5% BSA in PBS at RT and incubated with the primary antibody p-c-FOS (1:200, Cell Signaling, cat. 5348) or p-c-JUN (1:500, Cell Signaling, cat. 3270) in 0.5% BSA/PBS overnight at 4°C. The samples were thoroughly washed with 0.05% Tween-20/0.1% BSA in PBS whereafter the secondary antibody Alexa Fluor-488 (1:200, Thermo Fisher, cat. A-21206) was added in 0.5% BSA/PBS and incubated at RT for 2 hours. After washing, the samples were stained with DAPI (1:1000, Thermo Fisher, cat. 62248) in PBS and the chambers were removed and the slides were mounted using Prolong Gold (Invitrogen, cat. P36930). Images were acquired using the Leica SP8 (Leica, Wetzlar, Germany).

Paraffin-embedded tissue sections (5 µm thick) from mice hind limbs (ACVR1^Q207D/+^ or ACVR1^WT^)^28^ were prepared for immunofluorescence or Masson’s trichrome staining. Tissue sections were deparaffinized and rehydrated through a series of graded alcohols. For immunofluorescence, antigen retrieval was performed using a 10 mM sodium citrate buffer (pH 6.0) in a pressure cooker for 30 minutes. Afterwards, sections were washed and permeabilized with 0.2% Triton X-100/TBS supplemented with 1 mM CaCl_2_ and 1 mM MgCl_2_ for 20 minutes at RT. Next, sections were blocked using 5% BSA in TBST for 1 hour at RT followed by incubation with Alexa Fluor-488 conjugated wheat germ agglutinin (WGA) (1:250, Thermo Fisher, cat. W11261) for 30 minutes in the dark. After washing, a second blocking step with 5% BSA in TBST was performed for 1 hour. Primary antibody incubation was overnight at 4°C using c-FOS (1:2000, Cell Signaling, cat. 2250) or c-JUN (1:500, Cell Signaling, cat. 9165). Sections were then incubated with Alexa Fluor-555-conjugated secondary antibody (1:500, Thermo Fisher, cat. A42794) and co-stained with DAPI (1:1000, Thermo Fisher, cat. 62248) for 2 hours at RT followed by mounting. Immunofluorescent images were obtained using a Zeiss Apotome (Zeiss, Oberkochen, Germany). Masson’s trichrome staining was performed as previously described.^28^ Images were obtained using a Brightfield Eclipse E800 (Nikon, Tokyo, Japan).

### Osteochondrogenic differentiation, ALP and Alcian blue assays

Cells were seeded and cultured in the osteochondrogenic differentiation medium consisting of DMEM (Gibco, cat. 11965092) supplemented with 10 µL/mL Insulin-Transferrin-Selenium (ITS; Gibco, cat. 41400045), 200 µM Ascorbic Acid, 0.1 µM Dexamethasone, 350 µM L-proline, 1 mM Sodium Pyruvate, 1.25 µg/mL BSA and 100 U/mL P/S (differentiation medium; DM), which was refreshed twice a week. To test pharmacological intervention of ligand-induced differentiation, the DM was supplemented with Activin A (50 ng/ml), T-5224 (20 µM) or both vehicles (ligand buffer or DMSO) as experimental conditions for the entire length of the experiment. For the alkaline phosphatase (ALP) assays, hMSC- ALK2^WT^ or -ALK2^R206H^ were seeded at a density of 20^4 cells per well in a 48-wells format. The ALP staining was performed after 7 days of differentiation, while separately, the ALP activity was measured 11 days post-differentiation. 2D micromass cultures of hMSC-ALK2^WT^ or -ALK2^R206H^ were generated to determine chondrogenic matrix development, by seeding 30^5 cells in 10 µL droplets onto 24-wells plates and incubating for 2-3 hours prior to GM addition. The next day the micromass cultures were refreshed with differentiation medium as indicated in the figure. Differentiation was carried out for 3 weeks after which Alcian blue staining was performed.

For the ALP staining, the cells were washed with cold PBS and fixated using 3.7% formalin for 5 minutes at RT, washed with PBS and stained in ALP staining solution (2 mg NaASMX (Sigma), 6 mg Fast Blue (Sigma), 5 mL 0.2 M Tris (pH 8.9), 100 µL MgSO_4_ and dH_2_O up to 10 mL). After thorough washing, images were acquired using the Leica DMi8 with 10x magnification. Whole well densitometry from the total plate scan was performed using ImageJ (v1.53t). To measure ALP activity, the wells were washed twice with PBS, frozen at -80°C for 1 hour and lysed on ice for 1 hour using ALP lysis buffer (100 µM MgCl_2_, 10 µM ZnCl_2_, 10 mM Glycine (pH 10.5) and 0.1% Triton X-100). The active ALP was quantified by adding 20 µL lysate and 80 µL 6 mM PNPP (Pierce) in ALP lysis buffer to a clear 96-wells plate, incubating until the samples turned yellow and measured at an absorbance of 405 nm.

The 2D micromass cultures were stained by Alcian blue as described before.^29^ Images were acquired using the Leica DMi8 with 5x magnification. The staining was quantified by solubilization with 250 µL of 6M guanidine hydrochloride overnight at RT and subsequent absorbance measurements at 595 nm.

## Statistics

Statistical testing was performed using Graphpad Prism (v10) as indicated in every figure legend. P-values < 0.05 were considered statistically significant.

## 3. Results

### Phosphoproteomics identifies increased non-SMAD signaling including MAPK and RHO-associated GTPase pathways related to mechanotransduction in mutant ALK2^R206H^ expressing hMSCs

Dysregulated signaling in FOP extends beyond canonical SMAD pathways, but these non-canonical signaling networks remain poorly understood. To systematically map the altered signaling landscape in FOP, we performed phosphoproteomics of human MSCs overexpressing either wild-type ALK2 or the mutant ALK2^R206H^ (Figure 1A). As previously reported,^16^ our ALK2^R206H^-expressing cells showed aberrant SMAD1/5 phosphorylation in response to Activin A, confirming the expected gain-of-function phenotype (Figure 1B). Our phosphoproteomic analysis identified 9935 high-confidence phosphosites mapping to 2885 unique proteins, providing comprehensive coverage of the signaling networks in these cells (Supplementary Figure 1A-B). Similar to other studies, most phosphorylation events occurred on serine residues (∼90%), with threonine (∼9%) and tyrosine (∼1%) phosphorylation representing a minority.

**Figure 1.**
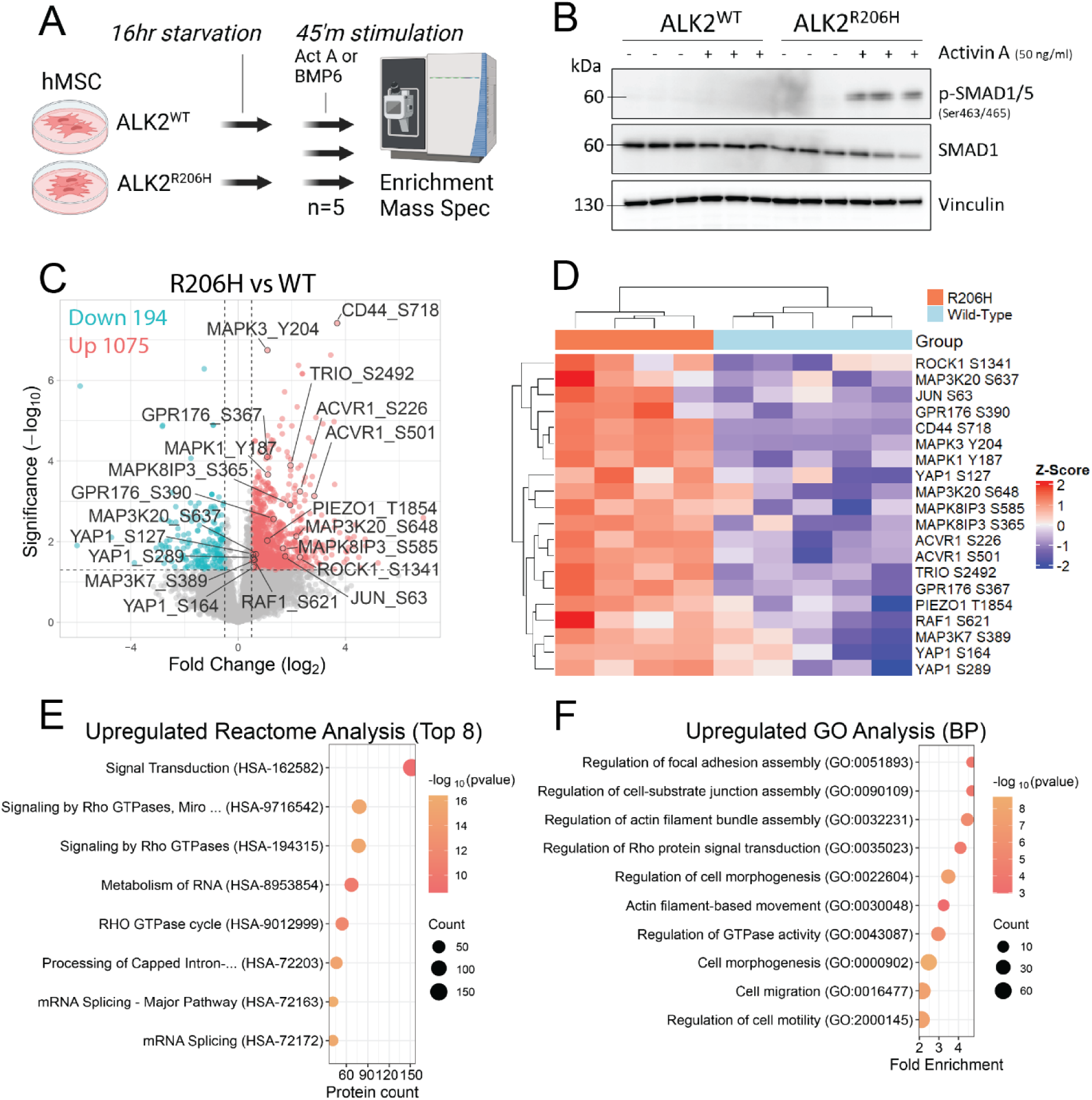
Phosphoproteomics reveals enhanced MAPK and RHO signaling, mechanosensing and cell morphogenesis in ALK2^R206H^ -expressing cells. A) Wild-type ALK2^WT^ and mutant ALK2^R206H^ expressing cells were serum starved for 16 hours, followed by stimulation with Activin A or BMP6 (both 50 ng/ml) for 45 minutes. B) Western blot analysis confirming aberrant SMAD1/5 phosphorylation in ALK2^R206H^ expressing cells upon Activin A stimulation. Three out of the five replicate samples used for mass spectrometry are shown. C) Volcano plot representing differential phosphorylation between unstimulated ALK2^R206H^ expressing cells and ALK2^WT^ controls. D) Heatmap showing the most interesting phosphosites comparing unstimulated ALK2^R206H^ and ALK2^WT^ expressing cells. E) Reactome pathway enrichment analysis of the differentially upregulated phosphorylated proteins identified in (C), highlighting the top 8 significantly upregulated pathways. F) Gene Ontology (biological process; BP) analysis of the upregulated hits from (C) reveals highly significant GO terms related to mechanosensing/transduction, cell motility and RHO-associated signal transduction.

Comparing unstimulated ALK2^R206H^ to ALK2^WT^ cells revealed 1269 significantly altered phosphosites, with the majority (1075 sites) showing increased phosphorylation in ALK2^R206H^ cells (Figure 1C). Among these, we identified previously unreported increased phosphorylation of ALK2^R206H^ at the highly conserved residues S226 and S501. Notably, we observed increased phosphorylation of multiple MAPK pathway components (MAPK1/ERK2, MAPK3/ERK1, MAPK8IP3/JIP3, RAF1) and mechanotransduction mediators (ROCK1, YAP1, PIEZO1) in ALK2^R206H^ cells (Figure 1D). Pathway analysis of the upregulated phosphoproteins showed significant enrichment of signal transduction networks, particularly RHO GTPase signaling and mRNA splicing pathways (Figure 1E). Gene Ontology analysis further revealed that the dysregulated proteins were predominantly involved in mechanosensing and cell morphogenesis (Figure 1F), suggesting that the p.R206H mutation in ALK2 fundamentally alters cellular mechanotransduction pathways.

To identify signaling events specifically triggered by ligand induction in FOP cells, we compared phosphoproteomic profiles upon Activin A or BMP6 stimulation. After analysing the ligand-induced hits, we determined that the most relevant players were identified downstream of Activin A stimulation, yielding a total of 145 upregulated ALK2^R206H^ specific Activin A-induced phosphosites not observed in ALK2^WT^ cells or under basal conditions (Figure 2A-B). These Activin A-specific changes included increased phosphorylation of key osteogenic regulators including mTOR (S1261), CTNNB1/β- catenin (S675), SOX6 (S98) and RUNX2 (S280).^39–41^ The 145 affected proteins were significantly enriched in cellular compartments associated with mechanosensing (Figure 2C), further supporting a central role for dysregulated mechanotransduction in FOP pathogenesis.

**Figure 2.**
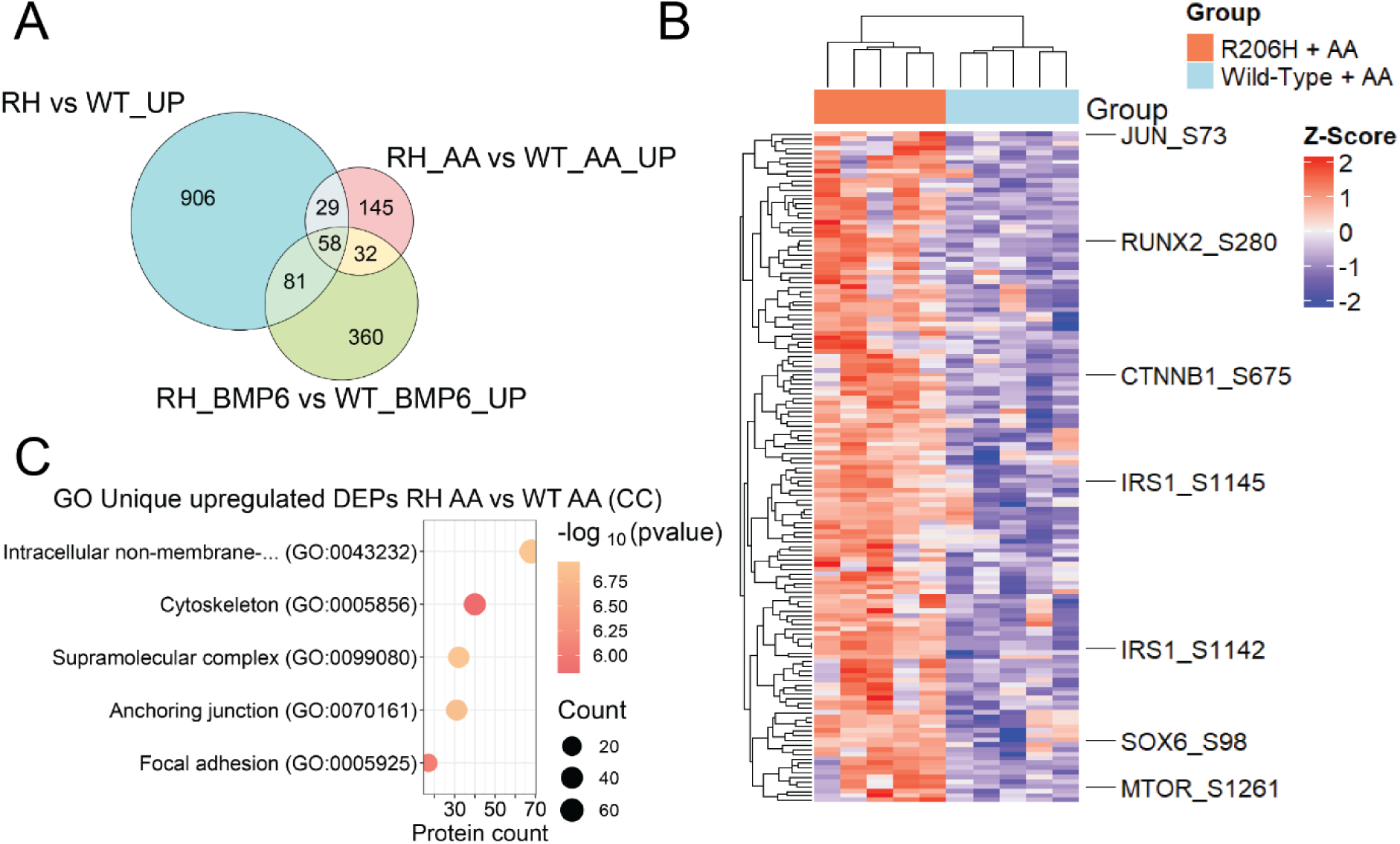
Activin A-induced non-SMAD signaling in ALK2^R206H^ cells involves mTOR, JUN and RUNX2 phosphorylation. A) Venn diagram representing differential phosphorylation analysis of significantly upregulated phosphosites in ALK2^R206H^ versus ALK2^WT^ conditions, comparing unstimulated (cian), BMP6 stimulated (green) and Activin A (AA) stimulated (red) cells. B) Heatmap of the 145 unique Activin A-induced upregulated phosphosites in ALK2^R206H^ showing annotation of the most interesting hits involving osteochondrogenesis. C) Gene Ontology analysis of the 145 upregulated phosphorylated proteins demonstrates that they can be allocated to cytoskeletal cellular compartments (CC) associated with mechano-sensing, such as anchoring junctions and focal adhesions.

### RNA sequencing confirms upregulated Activin A-induced RHO-mediated mechanotransduction and unveils increased Activator Protein-1 expression downstream of ALK2^R206H^ in FOP

To validate and extend these findings, we performed bulk RNA sequencing on ALK2^WT^ and ALK2^R206H^ expressing hMSCs with or without ligand stimulation (Figure 3A). Principal component analysis (PCA) showed that, while BMP6 induced similar transcriptional responses in both cell lines, Activin A triggered a unique transcriptional program specific to ALK2^R206H^-hMSCs (Figure 3B). These included the upregulation of canonical BMP target genes (*ID2-4*), mechanotransduction-associated factors (*KLF2/4/6, EGR1*) and RHO pathway targets (*CYR61*) (Figure 3D). Despite 16 h serum starvation, Gene Ontology analysis confirmed differential basal transcriptional profiles including increased cell motility, morphogenesis, and osteogenesis (involving osteoblast differentiation) in ALK2^R206H^ expressing cells (Figure 3C).

**Figure 3.**
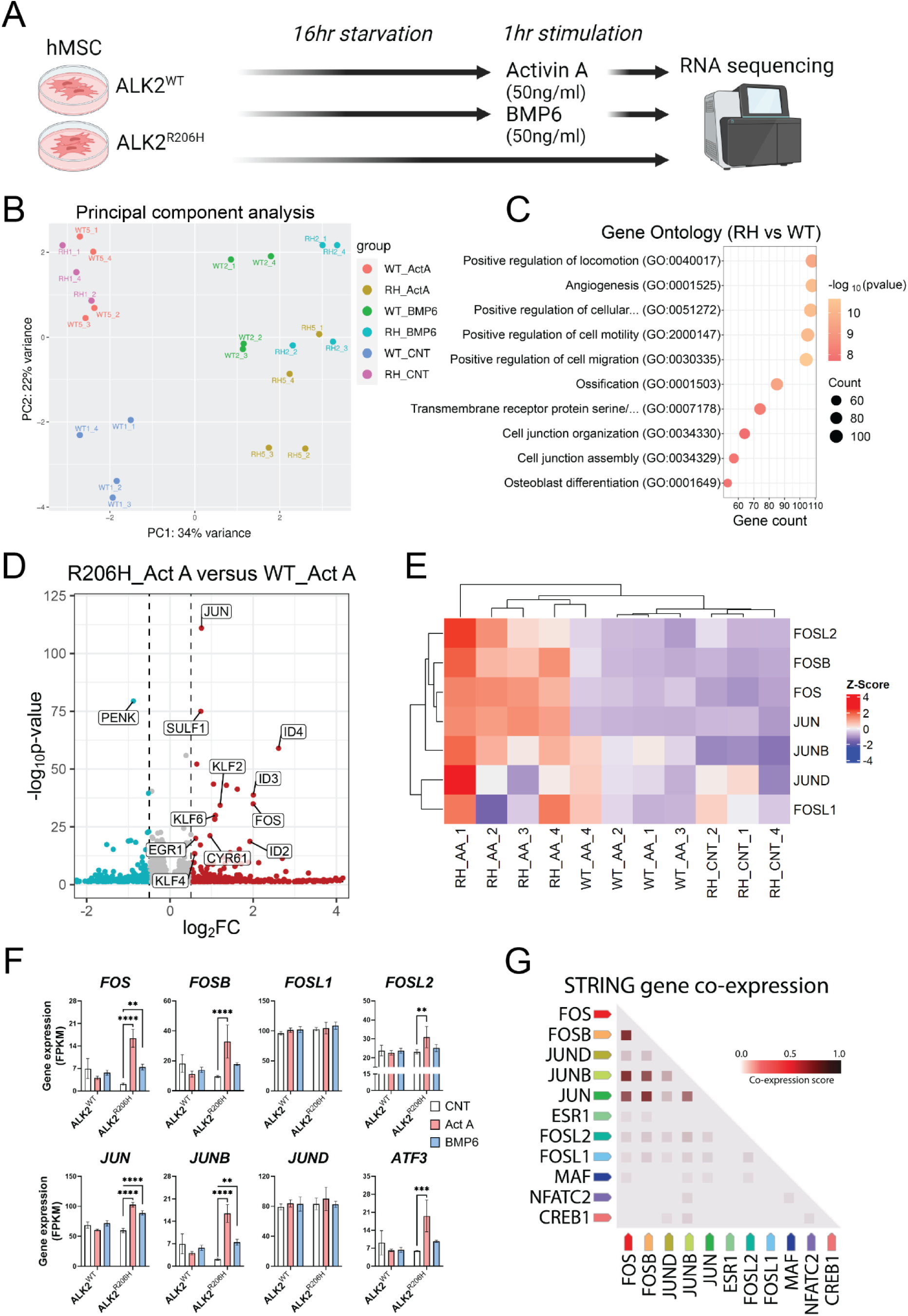
Transcriptomic analysis identifies upregulation of non-SMAD signaling, including mechanotransduction target genes like Activin A-induced Activator Protein-1 in ALK2^R206H^ hMSCs. A) Schematic representation of the experimental design prior to bulk RNA sequencing. hMSCs overexpressing ALK2^WT^ or ALK2^R206H^ were serum-starved for 16 hours and then stimulated with Activin A or BMP6 (50 ng/ml) for 1 hour before cell harvesting and RNA isolation. B) Principal component analysis (PCA) of all treatment groups showing clear clustering of all the replicates per group. Groups include ALK2^WT^ (WT) or ALK2^R206H^ (RH) cells treated with ligand buffer as control (CNT), Activin A (Act A, AA) or BMP6 (BMP6). C) Gene Ontology analysis showing the top 10 most significant terms based on differentially expressed genes (DEGs) comparing unstimulated ALK2^R206H^ cells to ALK2^WT^ controls. D) Volcano plot of the DEGs of ALK2^R206H^-cells treated with Activin A compared to its control group. E) Heatmap showing expression levels of all *FOS* and *JUN* family members highlighting significant overexpression of *FOS*, *FOSB*, *FOSL2*, *JUN*, and *JUNB* in ALK2^R206H^ cells following Activin A stimulation. F) Expression levels (in fragments per kilobase per million fragments; FPKM) of genes transcribing Activator Protein-1 (AP-1) factors in response to Activin A or BMP6 treatment. G) STRING co-expression analysis based on FOS expression. Statistical significance in (F) was tested using a two-way ANOVA following Šídák’s post-hoc tests: ** p<0.01, *** p<0.001, and **** p<0.0001.

Integration of the phosphoproteomics and transcriptomic data revealed a convergence on Activator Protein-1 (AP-1) transcription factors as important downstream mediators. As targets of mechanotransduction,^30,31^ regulators of inflammatory responses^32,33^ and drivers of osteochondrogenic homeostasis,^34^ these AP-1 family members may play a key role in HO. As such, multiple AP-1 family members, including *FOS, FOSB, FOSL2, JUN* and *JUNB,* were upregulated by Activin A stimulation in ALK2^R206H^ expressing hMSCs (Figure 3D-F). Interestingly, BMP6 also induced the expression of some AP- 1 factors (*FOS*, *FOSB*, and *JUN*) in ALK2^R206H^, although to a lesser extent than Activin A (Figure 3F). While tissue-wide co-expression analysis showed that these genes are transcribed ubiquitously (Figure 3G), our data demonstrates that ligand-induced AP-1 expression is specifically associated with ALK2^R206H^ signaling.

To validate these findings in disease-relevant contexts, we utilized FOP patient-derived iPSCs and FOP- like mouse models. Previously, we already developed and characterized one control and one FOP iPSC line generated from periodontal ligament fibroblast.^20^ Based on the latter, an isogenic control line (isoFOP) was established (Figure 4A).^35^ Further characterization using immunofluorescent staining and in vitro spontaneous trilineage differentiation confirmed the pluripotency of the isogenic line (Supplementary Figure 2). Control and FOP isogenic iPSC lines were differentiated towards the MSC lineage (iMSCs; Figure 4B). Similarly to our ALK2^R206H^ overexpression hMSCs lines, FOP iMSCs showed an induction of ALK2 downstream signaling in response to Activin A, as shown for SMAD1/5 phosphorylation and *ID1*/*ID3* gene expression (Figure 4C-E). Interestingly, basal *INHBA* expression levels were elevated in FOP iMSCs compared to its isogenic control (Figure 4E), previously reported for primary FOP skin fibroblasts^36^ and FOP iPSCs derived immune cells^37^, suggesting a detrimental feed forward loop. Importantly, like our hMSCs, these endogenous ALK2^R206H^ cells show Activin A-induced AP-1 expression not observed in isogenic control cells (Figure 4E).

**Figure 4.**
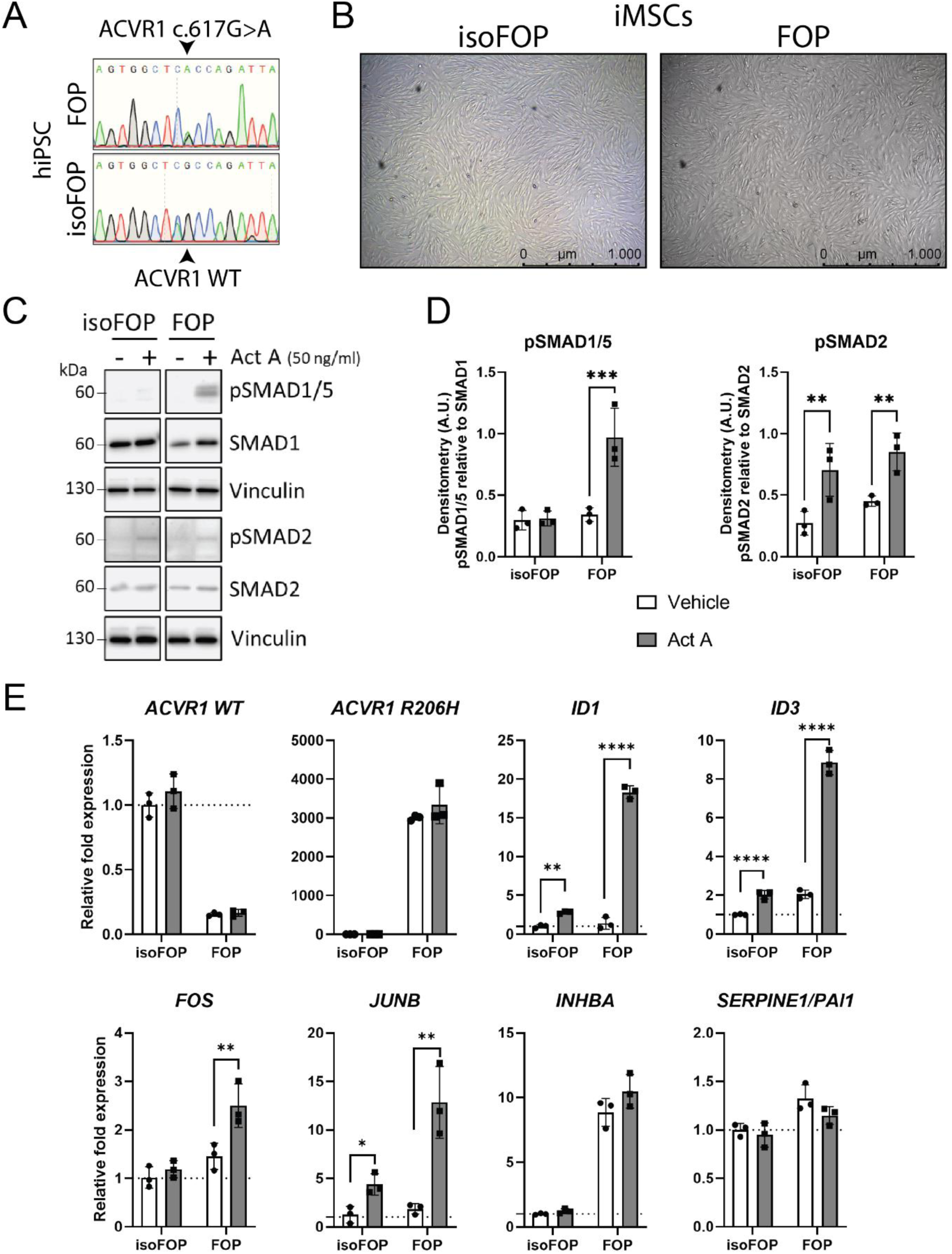
MSCs differentiated from newly established patient derived FOP and control hiPSC cells allows verification of Activin-induced AP-1 expression in FOP. A) Sanger sequencing of iPSC cells confirms the rescue of the heterozygous ACVR1 c.617G>A mutation, named the isogenic FOP line (isoFOP). B) Brightfield images of the FOP and isoFOP iMSCs shows a similar mesenchymal phenotype. C) Activin A signaling in isoFOP and FOP iMSCs assessed by western blotting. D) Quantification of the western blot in C). E) Gene expression analysis by RT-qPCR confirms expression of the mutant *ACVR1^R206H^* allele, consistent with enhanced *ID1/3* expression upon Activin A stimulation. FOP cells show substantial expression of *INHBA* compared to its isogenic control. Activin A stimulation in FOP derived iMSCs also induce AP-1 expression, shown by upregulated *FOS* and *JUNB* gene expression. Statistical significance tested by two-way ANOVA with Dunnett’s multiple comparison test: ** p<0.01, *** p<0.001, **** p<0.0001.

To confirm that AP-1 is also upregulated in vivo, we used the conditional ACVR1^Q207D/WT^ mice model to assess heterotopic ossification (HO) lesions. Immunofluorescence showed highly c-Jun and c-Fos rich areas within HO lesions (Figure 5), using consecutive tissue sections. Further, we found c-Fos and c-Jun positive mesenchymal cells within connective tissues of the transition zone of these HO lesions (data not shown). This implies that in vivo targeting of AP-1 might affect the HO lesions and the endochondral differentiation phenotype of connective/mesenchymal tissues in FOP.

**Figure 5.**
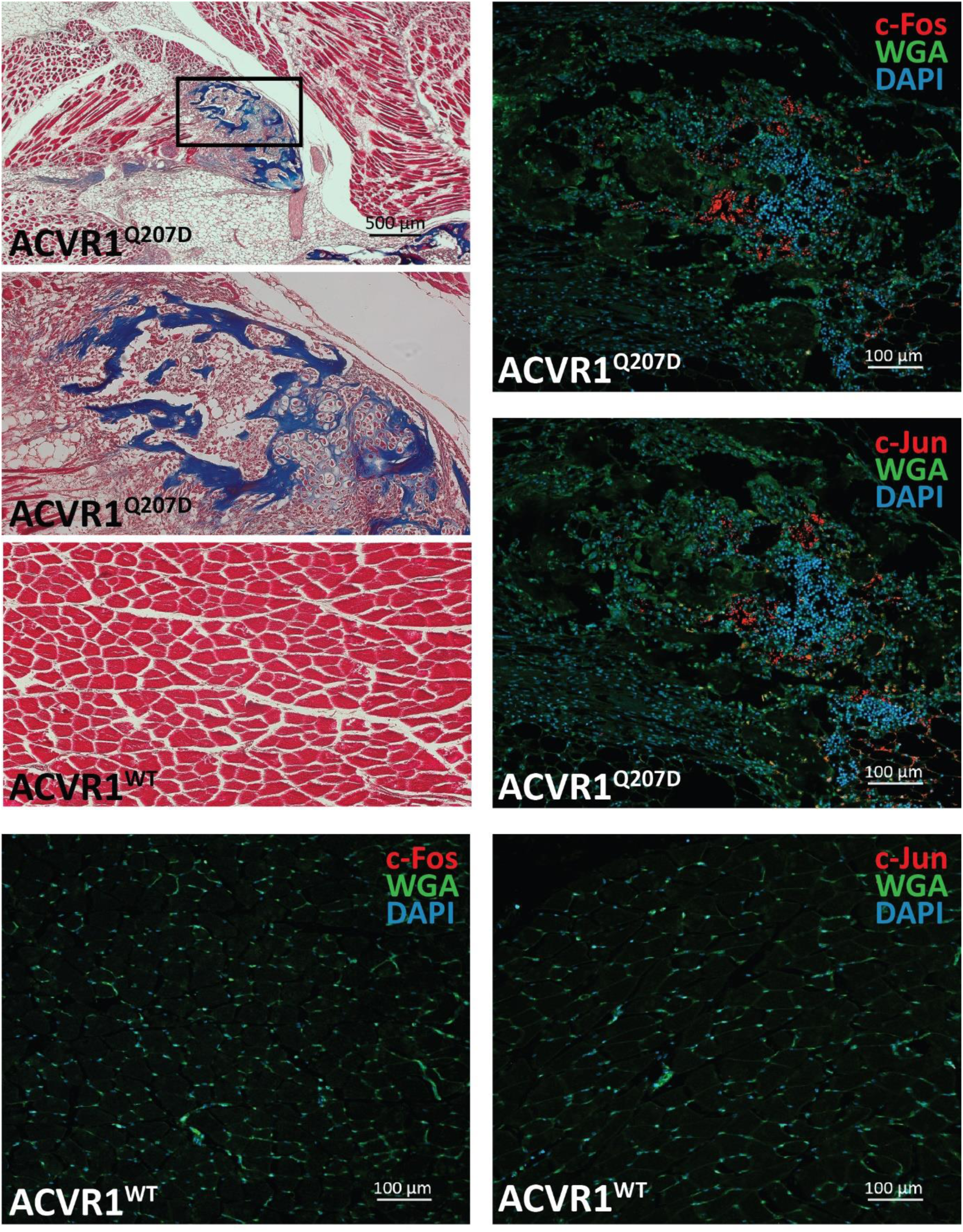
c-Fos and c-Jun proteins are abundantly present in HO lesions of ACVR1^Q207D/+^ mice. Histology and immunofluorescence analysis of an HO lesion of ACVR1^Q207D/+^ mice compared to muscle tissue of ACVR1^+/+^ mice. The Masson’s trichrome stain (upper left panels) shows muscle tissue in red and collagen rich sites in blue. Consecutive tissue sections were used for IF (other panels) of c-Fos or c-Jun with wheat germ agglutinin (WGA) and DAPI as a background stain to visualize cell membranes and nuclei, respectively.

### Activin A mediates context-dependent AP-1 activation and c-FOS phosphorylation through ALK2^R206H^

After identifying dysregulated AP-1 expression as a hallmark of ALK2^R206H^-mediated signaling, we next investigated its functional importance. Both phosphoproteomic and transcriptomic analysis suggested that AP-1 could be overactive in FOP. As such, c-JUN (Ser73) was phosphorylated (Figure 2B) and the previously identified AP-1 downstream targets *ADAMTS1* and *ID2*^26,38^ were upregulated by Activin A stimulation in our RNA sequencing data (Figure 3D). Based on these findings, we hypothesized that Activin A - ALK2^R206H^ signaling might directly involve AP-1 activation. To test this, we used conventional AP-1 responsive TRE-containing promoter reporters (like the *MMP1*-Luciferase) and phorbol 12- myristate-13-acetate (PMA) as positive control. We revealed context-dependent Activin A-induced AP- 1 activation, which was specific to ALK2^R206H^ signaling (Figure 6A-B). Importantly, this mechanism appears to be FOP and cell-type specific as ALK2^WT^ expressing cells did not respond to Activin A and non-human fibroblasts-like cells (COS-1) were incapable of TRE-dependent AP-1 activation. Binding of high affinity AP-1 complexes to the TRE is mainly mediated by c-FOS and c-JUN containing dimers.^26,39^ We hypothesize that c-FOS phosphorylation at Ser32 increases the active AP-1 dimers as Ser32 phosphorylation prevents protein degradation.^40,41^ To explore this, we used immunofluorescence to investigate if Activin A stimulation regulates c-FOS or c-JUN phosphorylation. We observed c-FOS phosphorylation at Ser32 in ALK2^R206H^ expressing hMSCs while c-JUN phosphorylation (Ser73) could not be validated (Figure 6C-F). These data suggest that, in addition to canonical SMAD1/5 activation, Activin A – ALK2^R206H^ signaling can activate non-SMAD pathways, which may play a role in modulating osteochondrogenic differentiation and maturation.^34^

**Figure 6.**
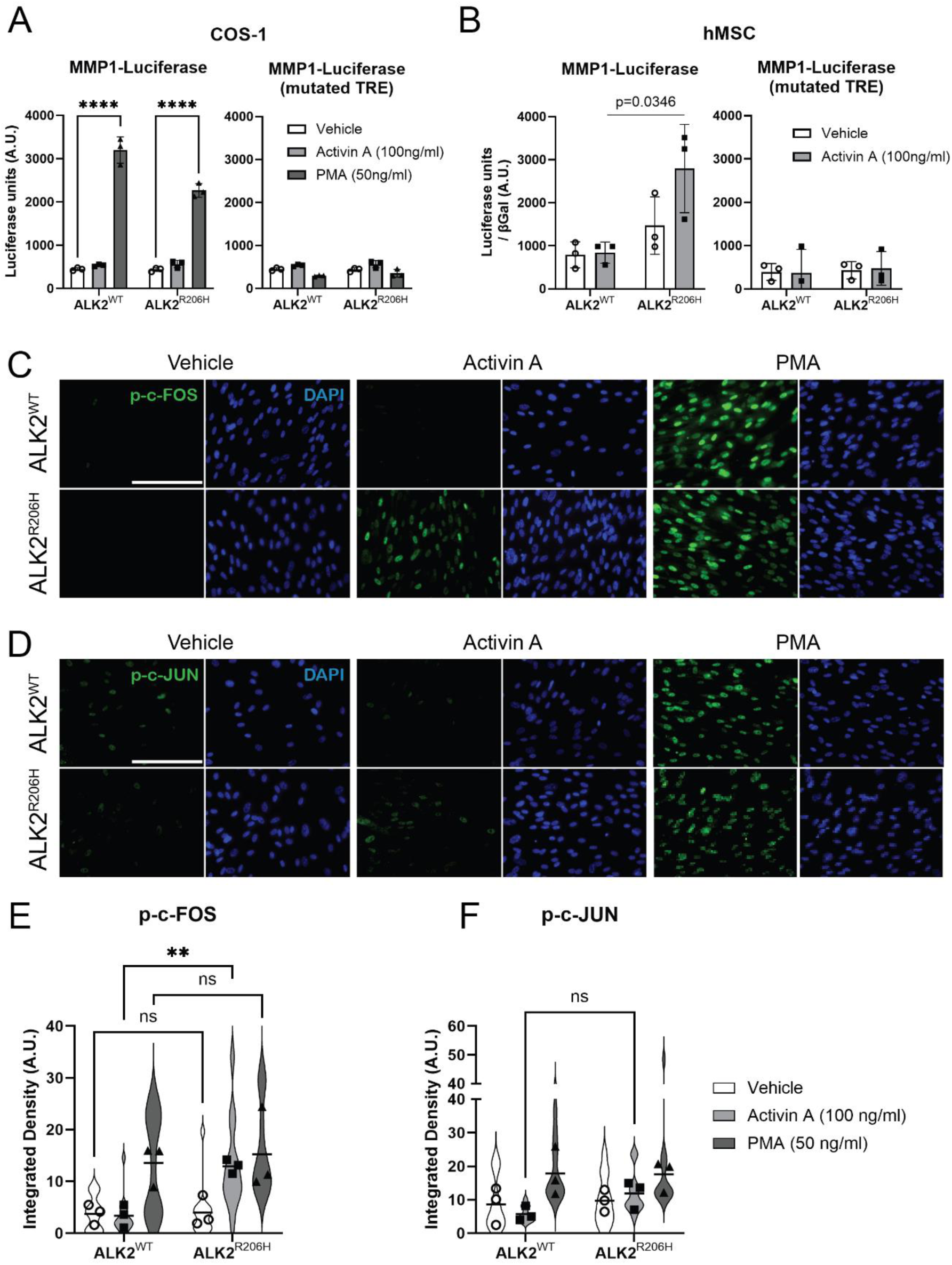
Context dependent TRE activation and FOS phosphorylation by Activin A signaling in ALK2^R206H^ expressing hMSCs. A) COS-1 cells co-transfected with an MMP1-promoter luciferase reporter construct containing or not (TRE) AP-1 DNA binding sites and ALK2 or ALK2^R206H^. Upon starvation, cells were stimulated with Activin A (100 ng/mL) for 48 hours. The phorbol agonist PMA is used as positive control. B) MMP1-reporter constructs were transfected into stably ALK2 or ALK2^R206H^ expressing human MSCs and stimulated with Activin A as indicated. C) Immunofluorescent staining of p-c-FOS (Ser32) on hMSCs expressing ALK2^WT^ or ALK2^R206H^. Scale bar represents 150 µm. D) Immunofluorescent staining of p-c-JUN (Ser73) on wild-type and mutant hMSCs. Scale bar represents 150 µm. E) and F) represent the quantification of C) and D), respectively. The mean of 4-5 images (violin plot) per independent replicate (data point) are depicted. Statistical significance tested by two-way ANOVA with Šídák’s in (A), Tukey’s in (B), (E) and (F) post-hoc tests: ** p<0.01, **** p<0.0001.

### Functional inhibition of AP-1 reduces osteochondrogenic differentiation in vitro

Based on the direct involvement of AP-1 in ALK2^R206H^-mediated signaling, we hypothesized that AP-1 could be a promising therapeutic target in FOP. Interestingly, genetic ablation of Fos/Jun family members reduced bone formation while transgenic overexpression of Fos/Jun increases bone formation in rodents.^34^ To explore whether functional inhibition of AP-1 could prevent Activin A-driven endochondral ossification in FOP, we used the experimental repurposed small molecule T-5224.^42^ Alkaline phosphatase (ALP) protein expression and activity, and Alcian blue staining on 2D micromass were assessed as surrogates for osteogenic and chondrogenic differentiation, both key processes in endochondral ossification. After 1 week of differentiation, T-5224 treatment completely blocked ALP expression in hMSCs expressing ALK2^WT^ and ALK2^R206H^ (Figure 7A-B). Similarly, we also observed normalization of the ALP activity when blocking AP-1 (Figure 7C). Interestingly, ALK2^R206H^ cells showed higher ALP expression and activity when cultured in differentiation medium compared to the ALK2^WT^ cells, irrespective of Activin A stimulation. This suggests that ALK2^R206H^ cells are inherently predisposed to osteochondrogenic differentiation, whereas wild-type cells require additional stimuli. After 3 weeks of culture, T-5224 treatment significantly reduced the formation of chondrogenic matrix, as shown by Alcian blue staining (Figure 7D-E). These findings suggest that increased AP-1 activation by Activin A in FOP osteochondroprogenitor cells is functionally relevant and therefore AP-1 inhibition may represent a novel druggable target for FOP to investigate in vivo.

**Figure 7.**
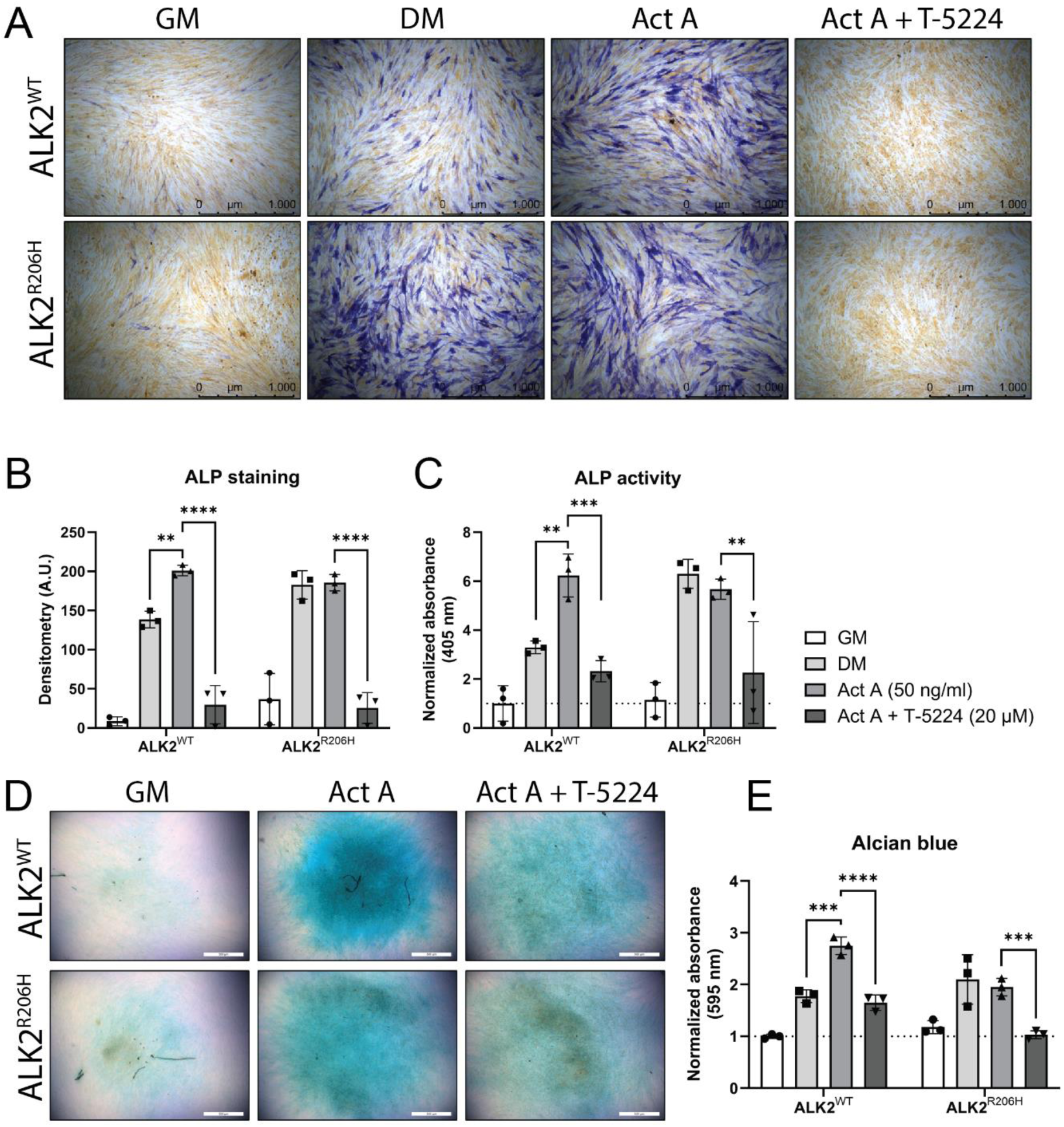
**Functional inhibition of AP-1 by T-5224 reduces osteochondrogenic differentiation in vitro.** A) ALP staining of hMSCs expressing ALK2^WT^ or ALK2^R206H^ cultured for 7 days in growth medium (GM), differentiation medium (DM), DM with Activin A (Act A; 50 ng/ml), or DM with Activin A and T-5224 (20 µM). Scale bar represents 1000 µm. B) Densitometric quantification of the ALP staining from triplicates. C) ALP activity assay measured after 11 days of culture under the indicated conditions. D) Alcian blue staining of 2D micromass cultures after 3 weeks of chondrogenic differentiation using the same treatment conditions as before: DM supplemented with Activin A (50 ng/ml) or Activin A and T-5224 (20 µM). Scale bar represents 500 µm. E) Quantified Alcian blue staining by absorbance measurements. Statistical significance was assessed using two-way ANOVA using Tukey’s multiple comparison tests: ** p<0.01, *** p<0.001, **** p<0.0001.

## 4. Discussion

In this study, we identified novel mediators of basal, BMP6-, and Activin A-induced ALK2^R206H^ signaling in FOP. Through phosphoproteomics, we characterized non-SMAD pathways downstream of ALK2^R206H^, including MAPK (e.g., ERK1/2, RAF1), RHO-associated pathways (e.g., ROCK1, YAP1, PIEZO1), and mTOR, along with RUNX2 activation. We showed that basal phosphorylation of ALK2 (Ser226 and Ser501) defines the mutant receptor. Bulk RNA sequencing confirmed transcriptional changes linked to mechanotransduction, highlighting the leucine zipper transcription factor complex Activator Protein-1 (AP-1) upregulation upon ALK2^R206H^ stimulation. These findings were validated in induced MSCs (iMSCs) derived from FOP patient iPSCs. Mechanistically, we also found that AP-1 is transcriptionally activated and c-FOS is phosphorylated in response to Activin A-ALK2^R206H^ signaling. Interestingly, high c-FOS and c-JUN proteins were detected in HO lesions in vivo. Finally, functional inhibition of AP-1 using the small molecule T-5224 successfully blocked osteochondrogenic differentiation in vitro.

Using our multi-omics approach, we identified dysregulated mTOR phosphorylation downstream of Activin A-ALK2^R206H^ in FOP. mTOR involvement in FOP has been previously demonstrated, with Rapamycin proposed as a therapeutic strategy now under clinical investigation in Japan.^17^ Similarly, our recent work showed that PI3Kα inhibition effectively blocks HO in pre-clinical models by reducing downstream mTOR-S6 activation.^16,28^ These findings underscore the potential of ALK2^R206H^-specific non-SMAD signaling pathways as novel druggable targets in FOP.

In line with the work of Shore and colleagues^43,44^, we identified activation of mechanotransductive mediators, including ROCK1, YAP1, and PIEZO1, in FOP. While their studies used embryonic fibroblasts from ALK2^R206H/+^ mice models, we confirmed similar pathway involvement in human MSC and patient-derived iMSC models. Phosphoproteomics and transcriptomics revealed activation of RHO-induced targets upon Activin A stimulation in FOP. These findings link disease-specific Activin A-induced ALK2^R206H^ signaling to mechanotransduction, highlighting its critical role in priming MSCs for endochondral differentiation.

For the first time, we identified AP-1, a downstream mechanotransduction complex^30,31^, as dysregulated in FOP. While AP-1 expression has been linked to TGF-β signaling, it has not been previously associated with BMP signaling.^45^ We show that Activin A-ALK2^R206H^ signaling induces AP-1 expression, likely independent of SMAD2/3 signaling, as suggested by the lack of *SERPINE1* expression upon Activin A stimulation. Additionally, BMP6 induces AP-1 expression in FOP cells but not in wild-type cells, suggesting the ALK2^R206H^-AP-1 axis operates via SMAD1/5-independent mechanisms or involves disease-specific non-SMAD co-factors. As such, we observed increased Activin A-induced RUNX2 phosphorylation, an osteochondrogenic transcription factor known to regulate *FOS* and *JUN*.^46^

A key question remains: which upstream non-SMAD mediator drives AP-1 expression and activation in FOP, including c-FOS phosphorylation at Ser32? The c-Jun N-terminal Kinase (JNK) is a promising candidate, as c-FOS phosphorylation at Ser32 has been shown to be JNK-dependent in cardiomyocytes treated with peroxide, independent of ERK, p38, or AKT signaling.^40^ Further, type I BMP receptors like ALK2 can activate JNK.^47,48^ Additionally, JNK activation occurs downstream of RHO/ROCK signaling^30^, which we observed to be elevated in our FOP model. Phosphorylation at Ser32 protects c-FOS from cytosolic ubiquitin-induced proteasomal degradation.^40,41^ It is also known that N-terminal c-JUN phosphorylation reduces its ability to form c-FOS/c-JUN dimers.^49^ While we did detect increased p-c-JUN (Ser73) through phosphoproteomics, we could not validate this through immunofluorescence. Combined with our MMP1-promoter luciferase data, this suggests the presence of an active c-FOS/c-JUN dimer in our FOP models.

AP-1 appears to be a promising target for FOP treatment, given its activation by mechanotransduction^20,21^, its pro-inflammatory function^22^, and its role in driving excessive bone conditions such as osteosarcoma and osteosclerosis.^34^ Previously AP-1 has been used to target the immune system in osteoarthritis, rheumatoid arthritis, and intervertebral disc degeneration with pre- clinical success.^27–29^ Also other drugs conventionally targeting arthritis are now being repurposed into FOP, like Tofacitinib and anti-IL1β.^61,62^ This triple-sided effect of AP-1 through osteochondrogenesis, mechanotransduction and inflammation is a good approach to target as HO is triggered by mechanical stress and flare-ups in FOP. Indeed, we showed the reduction of osteochondrogenic differentiation through AP-1 inhibition in vitro.

Currently, no pharmacological inhibitors targeting AP-1 have been approved for clinical use.^25,28^ In our study, we utilized T-5224 to block the DNA-binding activity of FOS-containing AP-1 complexes in vitro. T-5224 shows promise for FOP treatment due to its documented safety profile from a Phase 1 trial in Japan^27^, its immunosuppressive properties^29^, and its potential to inhibit the PI3K/AKT pathway.^63^ Notably, our previous research demonstrated that PI3Kα inhibition effectively blocks HO in FOP pre- clinical models^30,37^, further supporting the therapeutic potential of T-5224.

In conclusion, we have profiled multiple non-SMAD signaling pathways and found that the FOP-specific Activin A-induced ALK2^R206H^–c-FOS/AP-1-axis may be targeted to reduce osteochondrogenesis in FOP.

## 5. Author contributions

MW and GSD designed and conceptualized the study. FB and ALV were responsible for generating the iPSC-derived induced MSCs. SR, BS and PV performed the sample pretreatment, mass spectrometry and subsequent raw data analysis. CA and CF were responsible for the genetic rescue and characterization of the hiPSCs. MW generated all other experimental samples and performed all other experiments. MW performed all formal data analyses. GSD supervised the project. MJG, GSD and MW were responsible for funding acquisition. The original draft was written by MW. All authors critically revised the manuscript and MW finalized the article.

## Acknowledgements

Some of the figures were made using biorender.com using a license owned by GSD and MJG. Further, we acknowledge Hans van Dam for sharing the MMP1-Luciferase plasmids.

## 8. Data availability

The mass spectrometry proteomics data have been deposited to the ProteomeXchange Consortium via the PRIDE partner repository with the dataset identifier PXD058029. The RNA sequencing data have previously been deposited (GSE237512).^16^

---

**Supplementary Figure 1.**
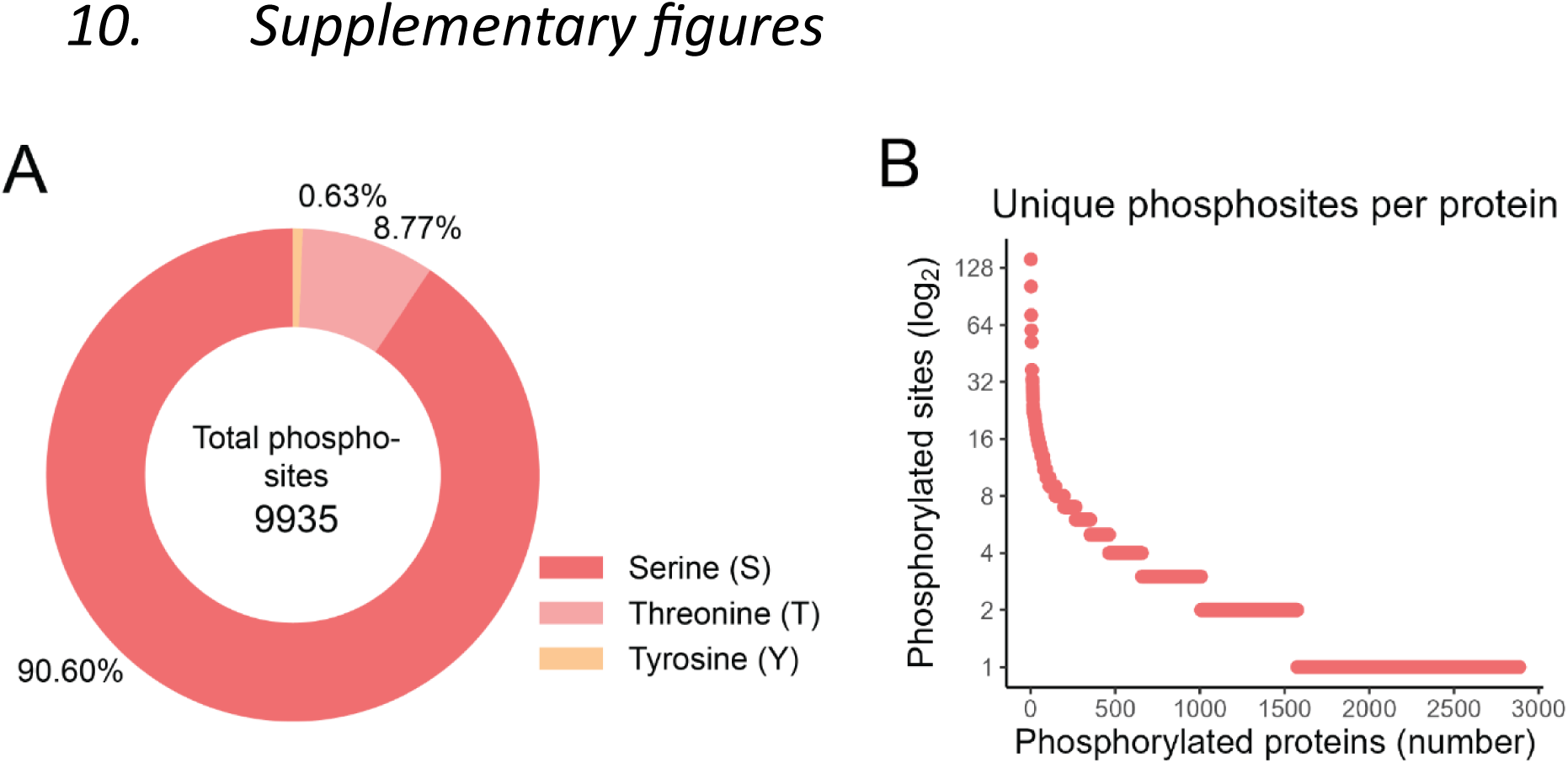
The phosphoproteomics experiment provides substantial depth for comprehensive signaling pathway analyses. A) Donut plot showing the total phosphosite yield (9935) stratified by the phosphorylated serine (S), threonine (T) or tyrosine (Y) amino acid residues. B) The unique phosphorylated sites can be mapped to 2885 different proteins with a decreasing phosphosite to protein distribution.

**Supplementary Figure 2.**
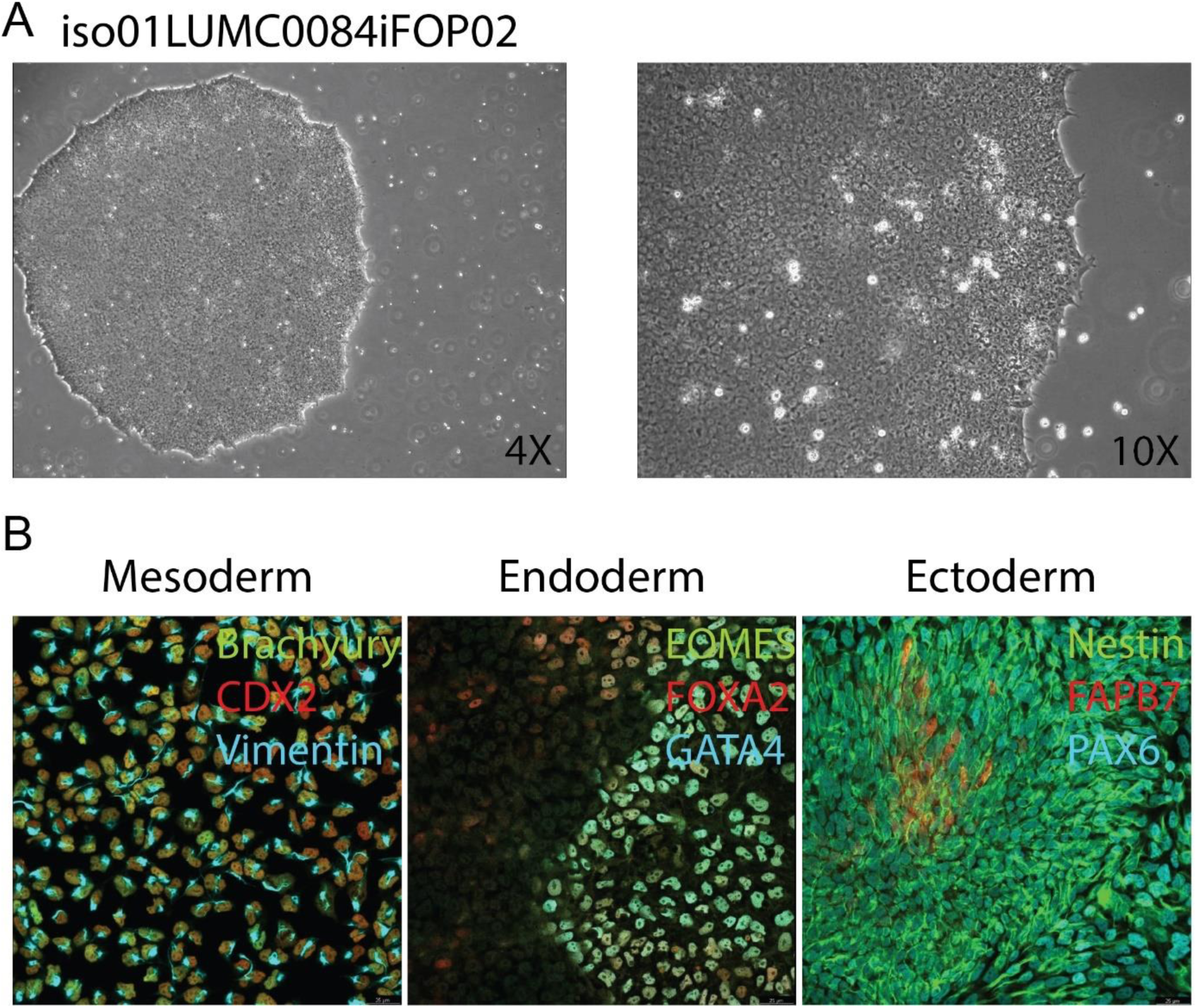
Validating pluripotency of the newly generated isogenic iPSC from the FOP patient-derived iPSC line. Brightfield and IF staining of the isogenic iPSCs to confirm pluripotency. A) Brightfield images of the isogenic line iso01LUMC0084iFOP02 derived from the FOP patient iPSC line 01LUMC0084iFOP02. Images are acquired using 4x or 10x magnification. B) After trilineage differentiation multiple lineage markers were stained; the mesoderm markers Brachyury, CDX2 and Vimentin, the endoderm markers EOMES, FOXA2, and GATA4, and the ectoderm markers Nestin, FAPB7, and PAX6. Scale bar represents 25 µm.

